# SAXS Curves of Detergent Micelles: Effects of Asymmetry, Shape Fluctuations, Disorder, and Atomic Details

**DOI:** 10.1101/815266

**Authors:** Miloš T. Ivanović, Markus R. Hermann, Maciej Wójcik, Javier Pérez, Jochen S. Hub

## Abstract

Small-angle X-ray scattering (SAXS) is a widely used experimental technique, providing structural and dynamic insight into soft-matter complexes and biomolecules under near-native conditions. However, interpreting the one-dimensional scattering profiles in terms of three-dimensional structures and ensembles remains challenging, partly because it is poorly understood how structural information is encoded along the measured scattering angle. We combined all-atom SAXS-restrained ensemble simulations, simplified continuum models, and SAXS experiments of a n-dodecyl-*β*-D-maltoside (DDM) micelle to decipher the effects of model asymmetry, shape fluctuations, atomic disorder, and atomic details on SAXS curves. Upon interpreting the small-angle regime, we find remarkable agreement between (i) a two-component tri-axial ellipsoid model fitted against the data with (ii) a SAXS-refined all-atom ensemble. However, continuum models fail at wider angles, even if they account for shape fluctuations, disorder, and asymmetry of the micelle. We conclude that modelling atomic details is mandatory for explaining SAXS curves at wider angles.

---

Detergent micelles are utilized in a wide spectrum of industrial, consumer, and scientific applications.^1,2^ For instance, because the cross-section of detergent micelles resembles lipid membranes,^3^ micelles are frequently used as lipid membrane mimics for solubilizing membrane proteins, thereby enabling further biophysical and structural studies.^4^ For a rational design of such protein-detergent complexes, and for modelling biophysical experiments, understanding of micellar shapes would be highly desirable, ^5,6^ with respect to both the overall shape and atomic details. However, owing to their intrinsic disorder and pronounced shape fluctuations, obtaining reliable models of micelles remains a major challenge.

Small-angle scattering, either with X-rays (SAXS) or neutrons, is a popular technique providing structural insight into soft-matter systems and biomolecules under near-native conditions.^7–17^ However, the interpretation of the one-dimensional scattering profiles in terms of structural models is challenging for several reasons: ^18^ (i) the information content of the SAXS profile is low and by far insufficient for defining all degrees of freedom of the solute, leading to a significant risk of overfitting the data; (ii) because the SAXS profile reports on the overall electron density contrast of the biomolecule, the data reflects the modulation of the solvent density in the hydration layer, suggesting that the hydration layer must be modelled upon interpreting the data. These challenges prompted the development of methods for the interpretation of SAXS data based on explicit-solvent molecular dynamics (MD) simulations because the simulations (i) add physicochemical information to the low-information SAXS data, thereby reducing the risk of overfitting the data, ^19^ and (ii) MD simulations may naturally account for the hydration layer of the solute.^20–23^

However, additional challenges emerge from a lack of understanding on how structural and dynamic information is encoded along the measured scattering angle. An exception is the very low-angle Guinier regime, which provides the radius of gyration of the solute.^24^ Further, for SAXS data of detergent micelles, the position of a broad maximum of the intensity curve *I*(*q*) was shown to correlate with the headgroup-headgroup distance across the shortest micelle diameter (see Fig. 1A/B),^12^ while the position of the first minimum was shown to be sensitive to the overall micelle volume.^25^ However, the information in the magnitude of the *I*(*q*) features, and in particular the information at wider scattering angles is poorly understood, which complicates the interpretation of the data. For instance, given that an experimental SAXS curve differs from a curve computed from a structural model, it is often unclear if such discrepancy originates from experimental problems or from a simplification in the model, such as an assumed model symmetry, neglect of shape fluctuations, or neglect of atomic details.

**Figure 1:**
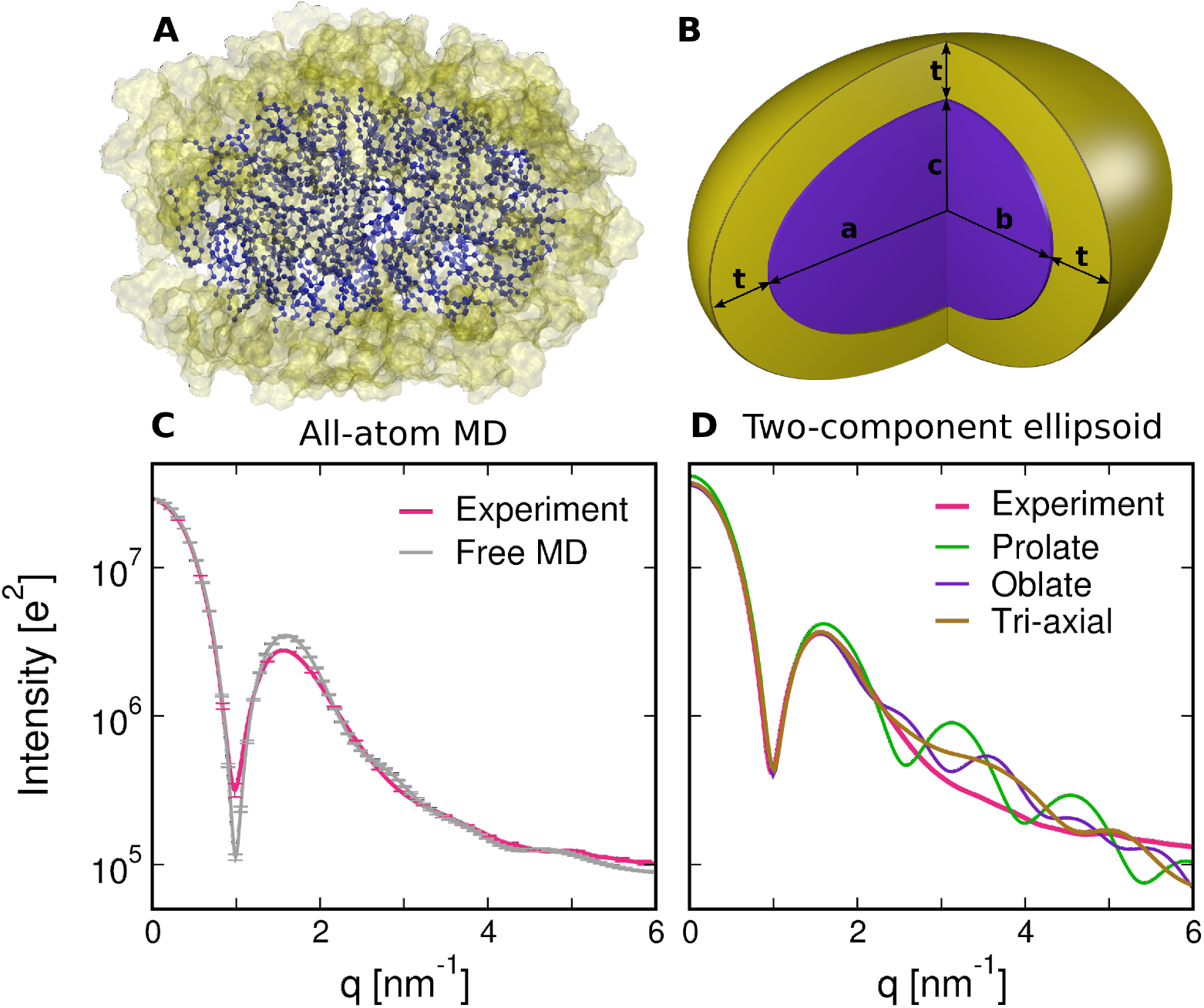
(A) Snapshot of a DDM micelle from all-atom MD simulation. The hydrophobic core is represented by violet spheres and sticks, and the hydrophilic headgroups are shown as yellow surface. Solvent is omitted for clarity. (B) Two-component ellipsoid model of a micelle. The same color scheme was used as in panel A. *a*, *b* and *c* denote the lengths of the semi-axes of the hydrophobic core. For *b* = *c* > *a*, the ellipsoid would be oblate; for *b* = *c* < *a* the ellipsoid would be prolate. Throughout this study, the thickness of the headgroup region was set to *t* = 0.55 nm (SI Methods). (C) Comparison of the experimental curve (red) with curves calculated from free, unbiased MD simulation (gray). For clarity, every 5th error bar is shown. (D) Comparison of the experimental SAXS curve (red) with the best-fitting curves computed from a two-component model: prolate model (green), oblate model (purple), and general tri-axial ellipsoid (*a* ≠ *b* ≠ *c*, brown).

To investigate the structural information in SAXS curves of soft-matter complexes, we measured the SAXS curve of a n-dodecyl-*β*-D-maltoside (DDM) detergent micelle^26^ up to *q* = 6 nm^−1^, where *q* = 4*π* sin(*θ*)*/λ* with the X-ray wavelength *λ* and the scattering angle 2*θ*. Using a recently developed method for coupling parallel-replica MD simulations to experimental SAXS data,^27^ we refined a heterogeneous atomic ensemble against the data with commitment to the principle of maximum entropy. ^28,29^ Having the atomistic ensemble in agreement with the data as a reference, we deciphered step-by-step the influence of model symmetry, shape fluctuations, disorder, and atomic detail on SAXS curve, by comparing the results from MD simulations with simplified micelle models. In addition, to shed more light on the complementarity of SAXS and MD, we investigated which *q*-range of the SAXS curve are most critical for improving the agreement of MD simulations with experimental conditions.

To obtain the atomic ensemble of the DDM micelle under experimental conditions, we used all-atom MD simulations. First, we used a series of free, unbiased MD simulations to determine the most likely aggregation number *N*_agg_, i.e. the number of detergent monomers per micelle. By comparing the position of the pronounced minimum at *q ≈* 1 nm^−1^ between the experimental curve and calculated curves,^25^ we found that the most likely *N*_agg_ under the given experimental conditions is 135 (Supporting Material). This value is in agreement with previously determined values at similar temperatures.^12,25,30^ In line with previous findings,^25^ the SAXS curve calculated from a free simulation of DDM micelle with *N*_agg_ monomers yield reasonable but not perfect agreement with the experiment, presumably as a consequence of minor imperfections of the applied CHARMM36 force-field^31^ (Fig. 1C).

Next, to overcome force-field imperfection, we refined the MD ensembles with an energetic restraint against the experimental curve. To apply only a minimal bias we ran several parallel replicas and coupled the replica-averaged SAXS curve to the experiment.^27^ This procedure follows Jaynes’ maximum entropy principle in the limit of a larger number of replicas, ^28,29^ and hence enforces that only the ensemble-averaged SAXS curve matches the experiment, but not necessarily the SAXS curve of each simulation frame. To test the effect of the number of parallel replicas, we refined ensembles using an increasing number of 1, 2, 4 (Movie S1), or 10 replicas, and we computed the SAXS curves and the micelle shape distributions from the refined heterogeneous ensembles (Fig. 2A/B, Table 1). In this work, the micelle shape was quantified via the semi-axes *a*, *b*, *c* of the hydrophobic core (see Fig. 1B), which were obtained from simulation frames every 10 ps via the instantaneous moments of inertia (SI Methods).

**Table 1:** Average semi-axes calculated from multi-replica SAXS-driven MD simulations (top rows) and from fitting a single two-component ellipsoid (bottom rows). Errors of MD simulations are given as 1 SEM of averages between independent runs. The prolate solution is unphysical because all semi-axes are significantly larger than the maximum extension of ~1.67 nm of the hydrophobic tail. ^32^

| All-atom MD<br># of replicas | a [nm] | b [nm] | c [nm] | t [nm] |
| --- | --- | --- | --- | --- |
| 1 | $3.29 \pm 0.02$ | $2.27 \pm 0.02$ | $1.68 \pm 0.02$ | |
| 2 | $3.17 \pm 0.03$ | $2.40 \pm 0.04$ | $1.67 \pm 0.01$ | |
| 4 | $3.16 \pm 0.01$ | $2.43 \pm 0.02$ | $1.63 \pm 0.01$ | |
| 10 | $3.17 \pm 0.03$ | $2.42 \pm 0.02$ | $1.67 \pm 0.01$ | |
| **Two-component model** |  |  |  |  |
| prolate (unphysical) | 3.39 | 1.97 | 1.97 | 0.57 |
| oblate | 2.84 | 2.84 | 1.60 | 0.55 |
| tri-axial | 3.20 | 2.47 | 1.65 | 0.55 |

**Figure 2:**
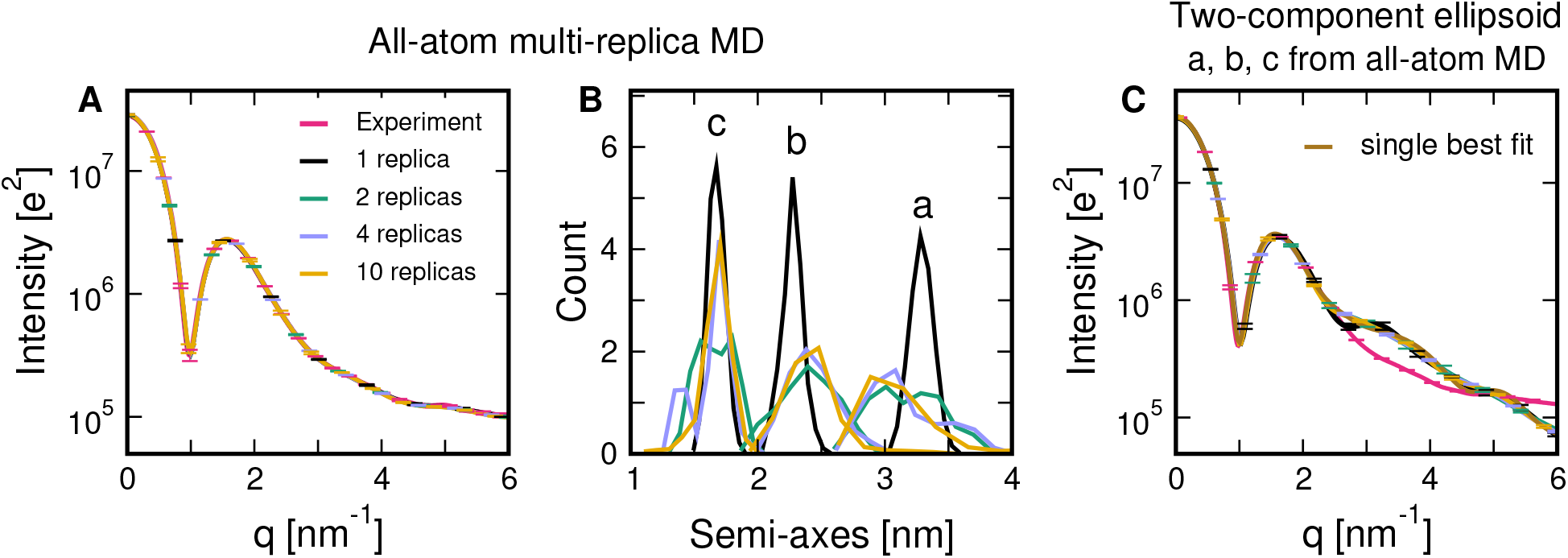
(A) Favorable agreement between the experimental SAXS curve (red) with curves from multi-replica SAXS-driven simulations refined against the experimental curve, shown for different numbers of parallel replicas (see legend). A few representative errors are shown as 1 SEM computed from independent runs. (B) Histograms of semi-axes *a*, *b*, *c* calculated from multi-replica SAXS-driven simulations obtained from the refined atomic ensembles. (C) Averages of curves calculated from the two-component tri-axial ellipsoid using semi-axes distributions from the refined atomic ensembles in panel (B). Representative errors show 1 SEM, computed via semi-axes distributions from independent MD simulation runs.

As expected, coupling only a single replica yields relatively narrow distributions of *a*, *b*, *c*, indicating an overly restrained ensemble (Fig. 2B, black) and a violation of the maximum entropy principle. Using multi-replica refinement, in contrast, the distributions become wider, in accordance to the maximum entropy principle (Fig. 2B, colored). The average semi-axes agree among multiple-replica simulations, and they differ by only ≈ 0.1 nm from the values obtained from the single-replica simulations. However, irrespective of the number of replicas, all refined ensembles reveal quantitative agreement with the experimental curve (Fig. 2A), suggesting that the SAXS curve encodes mainly the information about the mean micelle shape, and much less the information about the heterogeneity of the ensemble. Instead, MD simulations coupled to SAXS data with a minimal bias, as done here, are required to derive both the mean shape and the shape fluctuations.

Notably, in simulations with 10 or 20 parallel replicas, we reproducibly observed an unexpected horseshoe-shaped micelle in one or two replicas, respectively (Fig. S2). Although we cannot exclude that DDM micelles occasionally adopt elongated shapes, as reported for other detergent micelles,^10,33,34^ these shapes may indicate a force field limitation and hence may provide a starting point for further refinements of the CHARMM36 parameters (see SI Text).

The atomic ensembles of micelles in agreement with experimental SAXS data derived above provide a reference to study the influence of model symmetry, shape fluctuations, and atomic details on SAXS curves of detergent micelles. To this end, we investigated which parts of the SAXS curve may be explained with a greatly simplified two-component ellipsoidal micelle model, composed of uniform densities for head group and tail regions, as illustrated in Fig. 1B. Such models, constrained to oblate (*b* = *c* > *a*) or prolate (*b* = *c* < *a*) shapes, have well explained experimental curves of DDM micelles up to ~ 2.7 nm^−1^,^12,14,35,36^ and they were successfully applied to derive the aggregation number of micelles.^12,14,32^ Critically, fitting such models often leads to two disparate solutions, one prolate and one oblate, that match the data equally well.^37^ Since the existence of the water droplet or vacuum void in the micelle hydrophobic core would be energetically unfavourable, ^38^ the physically relevant solution was chosen by requesting that at least one semi-axis is shorter than the tail length, thereby avoiding a void at the micelle core. Following this procedure, and in line with previous findings, ^12^ we confirmed that both the oblate and prolate solutions fit the data well at small angles, where only the oblate solution avoids a vacuum void at the micelle core. At wide angles, however, where the experimental curve continuously decays along *q*, both the oblate and prolate solutions reveal several minima and maxima in sharp contrast to the data (Fig. 1D). Moreover, *a*, *b*, *c* determined with the oblate/prolate fits disagree with the values determined using SAXS-driven MD (Table 1).

The disagreement at wide angles may potentially be consequence of several simplifications: (i) the two-component oblate/prolate model allows for only two independent semiaxes, while the micelle under the experimental conditions most likely adopts the shape of a less symmetric, general tri-axial ellipsoid, as suggested by our previous study;^25^ (ii) by fitting a single model, shape fluctuations of the micelle in solution are ignored; (iii) two-component ellipsoid model assumes sharp core/headgroup and headgroup/water boundaries, while in reality these boundaries are more disordered and smeared out over a range of ~1 nm;^25^ (iv) atomic details of both the micelle and the solvent may have a significant effect on the SAXS curve at *q* > 2.5 nm^−1^. In the following, we disentangle the contribution of these potential sources of disagreement between model and experiment, with the aim to obtain an intuitive interpretation of the structural information of the wide-angle data.

First, to test the influence of the model asymmetry, we dropped the constraint to prolate/oblate shapes and instead fitted a two-component model of a general tri-axial ellipsoid to the experimental curve. The SAXS curve of the two-component tri-axial ellipsoid was computed following Ref. 39 (SI Methods), and the fits carried out by Powell optimization rapidly converged to a well-defined single optimum. The tri-axial ellipsoid fits the data only slightly better as compared to the prolate/oblate model (Fig. 1D brown). Specifically, the minima and maxima exhibited by the prolate and oblate models at *q* > 2.5 nm^−1^ are less pronounced in the case of a tri-axial ellipsoid reflecting the reduced symmetry. Nonetheless, the overall agreement to experiment at *q* > 2.5 nm^−1^ remains poor, suggesting that asymmetry is not the key to rationalize the wide-angle data. It is interesting to note, however, that the semi-axes *a*, *b*, *c* of the fitted tri-axial ellipsoid (i) were quite robust, irrespective of the fitted *q*-range, and (ii) favourably agree with the ensemble-refined MD simulations within 0.5 Å(Table 1). This finding suggests that the overall DDM micelle shape is well encoded in the *q* < 2.5 nm^−1^ range of the SAXS curve and may be extracted by fitting a tri-axial two-component model.

Second, to investigate the influence of the shape fluctuations on the SAXS curve, we generalized the two-component tri-axial ellipsoid model to a fluctuating model by averaging over a distribution of semi-axes *a*, *b*, *c*. Here, samples of the semi-axes were taken from snapshots of the multi-replica SAXS-driven MD simulations with 1, 2, 4, or 10 replicas; as such, the samples of semi-axes are compatible with the experimental conditions and, given a sufficient number of parallel replicas, reflect physically realistic shape fluctuations. However, including shape fluctuations into the two-component tri-axial model only marginally improves the agreement to experiment, as compared to a single best-fitting model (Fig. 2C). Namely, although the spurious bumps *q* > 2.5 nm^−1^ are partly smeared out, the calculated curves decay too rapidly with *q* as compared to experiment. This finding further confirms that the SAXS curve of DDM micelle is mainly given by the average micelle shape, whereas shape fluctuations have only the minor impact on the SAXS curve. Notably, this finding is not trivial because SAXS curves of other disordered ensembles, such as ensembles of intrinsically disordered proteins (IDPs), could not be explained by a single average structure.^40,41^ This difference is likely a consequence of the moderate magnitude the micelle fluctuations as compared to the large fluctuations carried out by many IDPs. Further, the fact that greatly different distributions of *a*, *b*, *c* (from a single set up to heterogeneous distributions) lead to nearly identical SAXS curves up to 6 nm^−1^ (Fig. 2C) implies that micelle fluctuations can not be derived from a SAXS experiment alone.

Third, to investigate the effect of disorder at the core-headgroup and headgroup–water interfaces, we smeared out the density contrast along the radial direction with a simple Gaussian filter, providing a more realistic density profile (Fig. 3, inset). The SAXS curve were computed analytically by generalizing the two-component model to a *N*-component model, following Ref. 39 (SI Methods). In line with the previous paragraph, shape fluctuations were included by averaging over semi-axes sampled from frames of the four-replica SAXS-driven MD simulation. However, upon smearing out the electron density profile, the agreement with the experimental SAXS curve becomes even worse, as apparent from an even more rapid decay of the calculated SAXS curve at *q* > 2.5 nm^−1^ (Fig. 3, green and red). The rapid decay of the SAXS curve of the *N*-component model (Fig. 3, green) may be rationalized by the loss of density–density correlations as a consequence a smeared out density.

**Figure 3:**
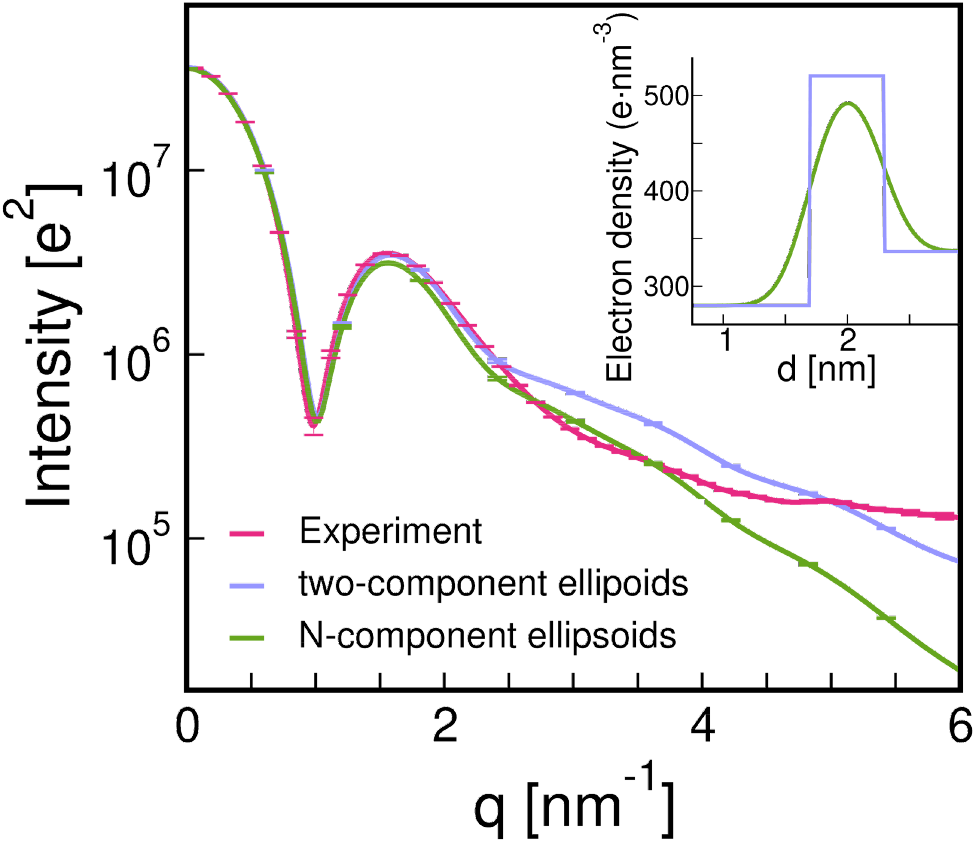
SAXS curves from experiment (red), the from a fluctuating two-component triaxial ellipsoid with a piecewise constant electron density profile (blue) and after smoothing the density profile with Gaussian filter (green). The inset shows an example for a piecewise constant and the smoothed electron density profiles along the minor micelle axis. The same samples of semi-axes were used as for the calculations for Fig. 2C. For clarity, only curves calculated from sets of semi-axes taken from the four-replica simulations are shown. Curves calculated with semi-axes from ten-replica simulations are nearly identical and shown in Fig. S4.

Taken together, the analysis demonstrates that accounting for shape asymmetry, fluctuations, and disorder may improve the agreement with experiment at wide angles only in a qualitative manner. Specifically, they largely remove the marked maxima and minima exhibited by the SAXS curve of the two-component oblate/prolate models, which indicated a spuriously high degree of symmetry and order in the oblate/prolate models (compare Fig. 1D with Fig. 3). To explain the wide-angle data quantitatively, we propose that, in addition, account for atomic details of both the micelle and the solvent are required, as captured by the MD simulations.

To decipher the role of atomic details on the SAXS curve at wider larger angles, we calculated the pair-distance distribution function *P* (*r*) from the calculated and from the experimental SAXS curves using the GNOM software^42^ (Fig. 4A and Fig. S5). The *P* (*r*) function is sensitive to density distribution of the micelle and provides more intuitive, real-space structural information. Evidently, the *P*(*r*) function from the ensemble-refined MD simulations favourably agree with the experiment (Fig. S5), as expected from the agreement of the SAXS curve. In contrast, the *P*(*r*) function obtained from the two-component models differ from the experiment, most prominently at small distances *r* between 0.9 nm and 3 nm. Namely, the features in the experimental or MD-based *P*(*r*) are smeared out by the two-component models, indicating a lack of structure at short-range, molecular distances (Fig. 4A). In addition, the *P*(*r*) from the prolate/oblate fits strongly differ at larger distances from the experimental *P*(*r*), reflecting a too high degree of symmetry as compared to the experiment (Fig. S5). To test how the lack of short-range structure propagates into the SAXS curve, we overwrote the *P*(*r*) from the shape-fluctuating two-component model with the experimental *P*(*r*) in different *r*-intervals, and subsequently back-calculated the SAXS curve via^43^ 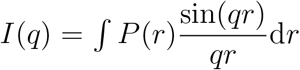, using the pddffit module of the ATSAS software (Fig. 4B).^44^ Although there is no simple one-by-one relation between specific *r*-regions of *P*(*r*) with *q*-regions of *I*(*q*), this analysis confirms that the short-range order (0 nm < *r* < 3 nm) has a strong effect on the SAXS curve at wider angles (*q* > 2.5 nm^−1^). Remarkebly, by replacing the region *r* = 0 nm to *r* = 3 nm of the the shape-fluctuating two-component model with the experimental *P*(*r*), we obtained very good agreement between experimental and calculated curve, suggesting that the discrepancy between experimental and SAXS curve calculated from fluctuating two-component ellipsoids is mainly recorded in the *r* = 0 nm to *r* = 3 nm region of the *P*(*r*).

**Figure 4:**
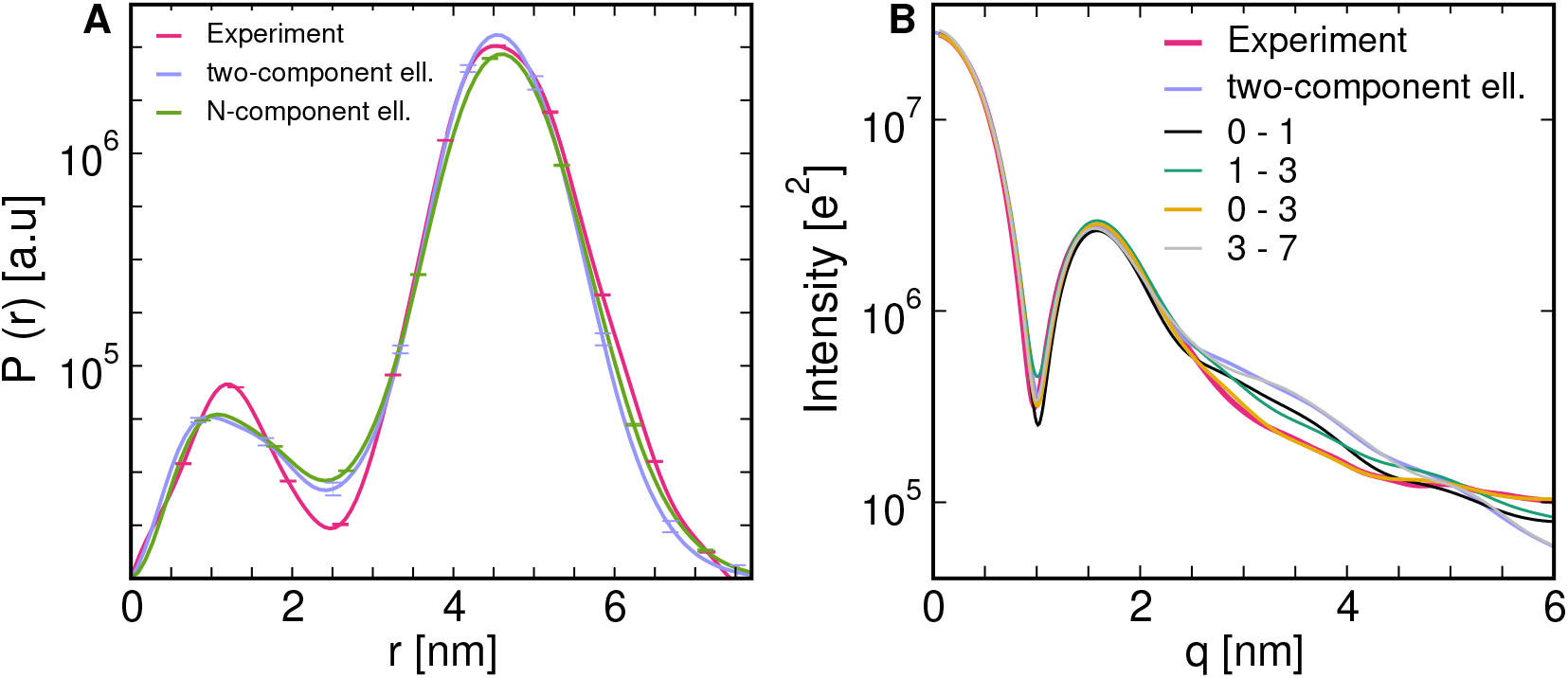
(A) *P*(*r*) functions calculated from the SAXS curves shown in (A) using the GNOM software.^42^ Every 10th error bar is shown, for clarity. *P*(*r*) curves calculated from the SAXS curves of oblate, prolate, or tri-axial fits and from the MD simulations are shown in Fig. S5 (B) To test the reason for discrepancy of *P*(*r*) functions calculated from experimental curve and shape-fluctuating two-component ellipsoids, we replaced parts of *P*(*r*) function of the shape-fluctuating two-component ellipsoids with the experimental *P*(*r*). Replaced *r*-ranges are shown in legend (units are all in nm).

To further investigate the structural information in different *q*-regions, and to test which part of the *q*-region of the SAXS curve plays the most important role in overcoming force-field imperfections during MD simulations, we performed the series of multi-replica SAXS-driven MD simulations using only specific *q*-intervals of the experimental curve as a target (Table 2 and Fig. S6). Table 2 lists the semi-axes of the refined micelles. The difference between the SAXS curves from experiment and from the refined MD ensembles were quantified with a non-weighted *χ*^2^ measure on a log scale, denoted 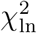. Computed SAXS curves (Fig. S6) as well as the calculated semi-axes show that: (i) applying the 0 nm^−1^ ≤ *q* ≤ 3 nm^−1^ region leads to results that are very close to results obtained with using the whole experimental curve. If instead even smaller-angle regions are applied (≤ 2 nm^−1^ or ≤ 1 nm^−1^), the agreement with the whole experimental curve still greatly improve. This finding demonstrates that the micelle shape is mainly encoded in the 0 nm^−1^ ≤ *q* ≤ 3 nm^−1^ region, as already indicated by fitting two-component tri-axial ellipsoid (see above). However, the Guinier region alone (≤ 0.5 nm^−1^) is insufficient for obtaining good agreement with the experiment; (ii) applying various intervals of the *q* > 3 nm^−1^ range leads only to a minor improvement compared to the free MD simulation. Taken together, these findings suggests that the MD force field already provides accurate description of the short-range order mainly encoded by the *q* > 3 nm^−1^ range, hence adding experimental data hardly improves the simulation. However, the force field alone has problems with defining the overall shape, as encoded by the *q* < 3 nm^−1^; hence experimental data in this range greatly improves the simulation.

**Table 2:** Semi-axes calculated from four- and ten-replica SAXS-driven simulations, using different *q*-intervals of the experimental curve as target curve. The deviation between calculated and experimental SA urves was quXaSntcified by a non-weighted *χ*^2^ on a logarithmic scale defined via 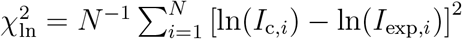, where *I*_*c,i*_ and *I*_exp,*i*_ denote the calculated and experimental SAXS intensities at the data point *i*, and *N* is the number of data points. Each four-replica (ten-replica) simulation was carried out for at least 100 ns (70 ns) per replica. Errors of *a*, *b*, *c* were computed using block averaging with 4 ns blocks, and errors were typically smaller than 0.02 nm. For reference, values from a free MD simulation are: *a* = 2.80 nm, *b* = 2.44 nm, *c* = 1.77 nm and 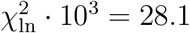

|  | 4 replicas |  |  |  | 10 replicas |  |  |  |
| --- | --- | --- | --- | --- | --- | --- | --- | --- |
| q-range | a [nm] | b [nm] | c [nm] | $\chi_{\ln}^2 \cdot 10^3$ | a [nm] | b [nm] | c [nm] | $\chi_{\ln}^2 \cdot 10^3$ |
| 0 - 6 | 3.15 | 2.44 | 1.63 | 0.5 | 3.17 | 2.42 | 1.67 | 0.6 |
| 0 - 4 | 3.17 | 2.43 | 1.65 | 0.8 | 3.19 | 2.39 | 1.68 | 0.8 |
| 0 - 3 | 3.18 | 2.40 | 1.64 | 1.0 | 3.22 | 2.36 | 1.67 | 1.1 |
| 0 - 2 | 3.13 | 2.48 | 1.61 | 5.3 | 3.15 | 2.43 | 1.62 | 4.7 |
| 0 - 1 | 3.17 | 2.26 | 1.70 | 6.3 | 3.19 | 2.25 | 1.72 | 7.1 |
| 0 - 0.5 | 2.85 | 2.48 | 1.73 | 23.2 | 2.81 | 2.47 | 1.76 | 28.9 |
| 1 - 2 | 3.09 | 2.45 | 1.62 | 5.3 | 3.13 | 2.44 | 1.62 | 3.9 |
| 2 - 4 | 3.13 | 2.40 | 1.69 | 1.8 | 3.05 | 2.41 | 1.68 | 3.0 |
| 2 - 3 | 3.08 | 2.42 | 1.68 | 2.5 | 3.05 | 2.43 | 1.67 | 4.3 |
| 3 - 4 | 2.96 | 2.38 | 1.75 | 18.5 | 2.89 | 2.44 | 1.73 | 21.5 |
| 3 - 6 | 2.94 | 2.41 | 1.73 | 24.0 | 2.93 | 2.42 | 1.73 | 18.9 |
| 4 - 6 | 2.86 | 2.41 | 1.78 | 26.9 | 2.87 | 2.42 | 1.76 | 27.8 |

To conclude, we obtained an heterogeneous atomic ensemble of a DDM detergent micelle by coupling a set of parallel-replica MD simulations to an experimental SAXS curve. Because the multi-replica ensemble refinement method applies only a minimal bias, as requested by Jaynes’ maximum entropy principle, the shape fluctuations of the free simulations were maintained. We found that scattering data at small angles (*q* < 3 nm^−1^) may guide the simulation into quantitative agreement with experiment, whereas scattering data at wider angles is matched already by free simulations with reasonable accuracy. This suggests that the force field is capable of reproducing the short-range structure of the micelle at atomic and molecular scales, but experimental data is needed to obtain the correct overall shape. According to the refined ensemble, the DDM micelle at 15°C adopts on average the shape of a general tri-axial ellipsoid. The major and middle semi-axes fluctuate by ~20% and the minor semi-axis by 5–10%.

The refined atomic ensemble provided a reference to test whether the fitting of simplified analytic models to the data may provide physically correct micellar shapes. Remarkably, by fitting a two-component general tri-axial ellipsoid to the data, we obtained a micellar shape in quantitative agreement with the multi-replica ensemble refinement, suggesting that (i) the two-component tri-axial model was not overfitted, and (ii) that the SAXS curve up to *q* ≈ 2.5 nm^−1^, in the case of DDM micelles, contains sufficient information for defining three independent semi-axis as well as the headgroup thickness. Upon restricting the fit to prolate or oblate shapes, however, we obtained different semi-axes, and the long-range structure quantified by the *P*(*r*) function disagreed with the experiment. Further, by increasing the complexity of the analytic micelle model step by step, we analyzed the role of model asymmetry, shape fluctuations, and disorder on the SAXS curve of the micelle. We found that these features partly improve the agreement with the experiment at wider angles, but, even when combined, they are insufficient for obtaining quantitative agreement. Taken together, atomic and molecular details, as naturally included in the MD simulation, are required to quantitatively explain the SAXS curve over the entire *q*-range.

## Supporting information

Supporting Material for: SAXS Curves of Detergent Micelles: Effects of Asymmetry, Shape Fluctuations, Disorder, and Atomic Details

## Acknowledgement

M.T.I., M.R.H. and J.S.H. were supported by the Deutsche Forschungsgemeinschaft (DFG) (HU 1971/3-1 and HU 1971/4-1). M.T.I. was additionally supported by the DFG via SFB 803/A12.

