## Supporting Material for: SAXS Curves of Detergent Micelles: Effects of Asymmetry, Shape Fluctuations, Disorder, and Atomic Details for "SAXS Curves of Detergent Micelles: Effects of Asymmetry, Shape Fluctuations, Disorder, and Atomic Details"


### Methods

#### SAXS experiment

The SAXS curve of the DDM micelle was taken from a recent study.[1] Details of sample preparation and data collection are provided in Ref. 1. For the purpose of this work, the experimental SAXS curve was smoothed using the default options of GNOM software.[2]

#### SAXS curves for tri-axial ellipsoid models

Detergent micelles have previously been modeled as two-component ellipsoids of revolution, i.e. ellipsoids with two density contrasts for tails and headgroups, and with only two independent semi-axes (prolate/oblate ellipsoids).[3] These calculations can be generalised to a general tri-axial ellipsoid, and/or to models composed of $N$ instead of two concentric shells. Here, the $N$ shells are constructed by superimposing $N$ ellipsoids. Each of the $N$ ellipsoids is defined by three semi-axes $a_i$, $b_i$ and $c_i$, the volume $V_i = \frac{4}{3}\pi a_i b_i c_i$, and by the solvent-subtracted electron density $\rho_i$, where $i = 1, \ldots, N$. The semi-axes are denoted in decreasing order, for instance $a_1 > a_2 > \ldots > a_N$. Following the Eqs. 55, 57, and 62 of Ref. 4, the scattering intensity of the $N$-component tri-axial ellipsoid is given by:

$$I(q, a_1, \ldots, a_N, b_1, \ldots, b_N, c_1, \ldots, c_N) = \frac{2}{\pi} \int_0^{\pi/2} \int_0^{\pi/2} F_3^2(q, r_1, \ldots, r_N) \sin\alpha \, \mathrm{d}\alpha \, \mathrm{d}\beta \tag{S1}$$

where

$$r_i(a_i, b_i, c_i, \alpha, \beta) = \left[ \left( a_i^2 \sin^2\beta + b_i^2 \cos^2\beta \right) \sin^2\alpha + c_i^2 \cos^2\alpha \right]^{1/2}. \tag{S2}$$

Here,

$$F_3(q, r_1, \ldots, r_N) = \rho_1 V_1 F_1(q, r_1) + \sum_{i=2}^{i=N} (\rho_i - \rho_{i-1}) V_i F_1(q, r_i) \tag{S3}$$

is the scattering amplitude of an $N$-shell ellipsoid, and

$$F_1(q, r_i) = \frac{3\left[\sin(qr_i) - qr_i \cos(qr_i)\right]}{(qr_i)^3}. \tag{S4}$$

For the two-component tri-axial ellipsoid (with only two shells), equation S3 simplifies to

$$\tilde{F_3} = \rho_1 V_1 F_1[q, r(a+t, b+t, c+t, \alpha, \beta)] + (\rho_2 - \rho_1) V_2 F_1[q, r(a, b, c, \alpha, \beta)], \tag{S5}$$

Here, the hydrophobic core is described as a tri-axial ellipsoid with semi-axes $a$, $b$, $c$ (Fig. 1B, purple region) with electron density $\rho_{\text{core}}$. The headgroup region has a constant thickness $t$ along the three axes (Fig. 1B, yellow region) and electron density $\rho_{\text{hg}}$. Accordingly, the parameters in Eq. S5 are given by $\rho_1 = \rho_{\text{hg}} - \rho_{\text{sol}}$, $\rho_2 = \rho_{\text{core}} - \rho_{\text{sol}}$, $V_1 = \frac{4}{3}\pi(a+t)(b+t)(c+t)$, and $V_2 = \frac{4}{3}\pi abc$. We validated numerically and analytically that the mathematical expressions shown above reduce to the expression for a two-component ellipsoid of revolution (oblate/prolate) used by Lipfert *et al.*[3]

The computed scattering intensity is sensitive to the electron densities, suggesting that reasonably accurate electron density estimates are required. For the density of the hydrophobic core and the headgroup region at the temperature of 15°C we used $\rho_{\text{core}} = 279.8\,\text{e}\,\text{nm}^{-3}$ and $\rho_{\text{hg}} = 520.5\,\text{e}\,\text{nm}^{-3}$. These values were determined taking into the account that: (i) electron densities of the core and headgroup region at 25°C are $277\,\text{e}\,\text{nm}^{-3}$ and $520\,\text{e}\,\text{nm}^{-3}$, respectively;[3] (ii) the density of the DDM detergent at 15°C is increased by 0.56% compared to 25°C;[5] (iii) the density of the alkyl tails at 15°C is increased by 1.02% compared to 25°C, as estimated by the temperature dependence of the density of alkanes with similar chain length.[6] The solvent density was set to $\rho_{\text{sol}} = 336.7\,\text{e}\,\text{nm}^{-3}$ to match the electron density of the 150 mM NaCl aqueous solution at the temperature of 15°C.

The thickness of the headgroup region, $t$, was determined once to 0.55 nm by fitting the two-component genereal tri-axial ellipsoid to the date. This value is close to the value of $t = 0.6$ nm, as previously determined by fitting a ellipsoid of revolution (prolate/oblate) at the temperature of 25°C.[3] It was shown previously that modulating $t$ in the range of 0.6 nm to 0.63 nm or using different headgroup thicknesses hardly influence the fits to the experimental data in the case of two-axial ellipsoids.[3] Likewise, we found here that small modulations of $t$ along the three axes hardly influences the fits in the case of a general tri-axial ellipsoid and hardly influences the results from modeling shape fluctuation. Therefore, in all follow-up calculations with the ellipsoid model, the value of $t = 0.55$ nm was used.

The smeared out electron density was modeled using an $N$-component tri-axial ellipsoid, where $N = 200$ was used. The density profiles along the three axes were obtained by convoluting the piecewise constant density (corresponding to a two-component ellipsoid) with a Gaussian filter with $\sigma = 0.2$ nm (inset in Fig. 3 A). Using this filter, the smoothed density profiles were qualitatively similar to the density profiles determined from MD simulations.[5] The largest semi-axes of the $N$-component tri-axial ellipsoid was chosen such that Gaussian tails up to $3\sigma$ of the smoothed density were taken into account.

#### MD simulations and SAXS calculations

**MD setup and simulation parameters** Unbiased, free simulations were carried out similar to previous work.[5] In short, detergent, water and ion interactions were modeled using CHARMM36 lipid force-field[7] and CHARMM-modified TIP3P water.[8] Free simulations were carried out with GROMACS 2018.3.[9] If not stated otherwise, the micelle was solvated in a 150 mM NaCl aqueous solution.[10] Likewise, a 150 mM NaCl solution was simulated as pure-solvent system. The temperature was controlled at 15°C to match the experimental conditions using velocity rescaling[11] ($\tau = 1$ ps). The pressure was controlled at 1 bar using the Berendsen barostat[12] ($\tau = 5$ ps). Electrostatic interactions were calculated using the particle-mesh Ewald method.[13,14] Dispersive interactions and short-range repulsion were described together by a Lennard-Jones potential with a cutoff at 1.2 nm.. Length of the free simulations, with and without added salt were 500 ns. The convergence of the free simulation ensemble was validated by comparing SAXS curves computed from 50-nanosecond blocks of the trajectory. The first 50 ns of all free simulations were removed for equilibration.

**Explicit-solvent SAXS curve predictions** SAXS curves were calculated using explicit-solvent SAXS predictions described previously.[15] Explicit solvent atoms contributing to the SAXS curve were defined by a spatial envelope. Here, the envelope was constructed at a distance of at least 1 nm from all detergent atoms in all frames of an equilibrium simulation. Because the micelle heavily fluctuates, this procedure led to the distance of ~2 nm between micelle and envelope in most frames. The same envelope was used for all SAXS-driven simulations. The SAXS curve was calculated using the positions of atoms inside the envelope each 10 ps. Scattering amplitudes were computed using 1200 **q**-vectors per $q$-point, which were distributed by the spiral method. Because the TIP3P solvent density differs from the experimental value, we corrected the solvent density to $335.7\,\mathrm{e\,nm^{-3}}$, corresponding to a 150mM NaCl solution, following the procedure described previously.[15]

**SAXS-driven MD simulations** The initial configurations of SAXS-driven simulations was taken from free simulations. In the case of parallel-replica simulations, frames for the replicas were taken from free MD snapshots at 5-nanosecond intervals. Apart from using 150 mM NaCl aqueous solution instead of pure-water solution, single-replica SAXS-driven simulations[16] were performed as described previously.[5] Details of the recently developed multi-replica SAXS-driven simulations, following the principle of maximum entropy, are described in Ref. 17. In all single-replica and parallel-replica simulations, the temperature was controlled at 15°C using a stochastic dynamics integrator ($\tau = 0.2\,\mathrm{ps}$). To validate that the refined ensembles are reproducible, we ran multiple independent simulations for each setup of 1, 2, 4, 10 or 20 replicas. The number of independent runs, the applied force constant used to couple the simulation to the data, and total simulation times are listed in Table S1. Further, we excluded that the memory time $\tau$ used for on-the-fly averaging of the SAXS curve[16] has a significant effect on the refined ensembles. We found that the choice for $\tau$ between 100 ps and 500 ps did not influence the calculated SAXS curve or semi-axes. Here, for all the production runs, we used $\tau = 200\,\mathrm{ps}$. During the first 5 ns of SAXS-driven simulations, the SAXS-derived forces were gradually switched on, and the first 8 ns of all SAXS-driven simulations were omitted from the analysis for equilibration. In contrast to previous work,[5] the overall scale $f$ and a constant offset $c$ of the experimental curve were marginalized out on-the-fly during the SAXS-driven simulations. As shown previously,[18] marginalizing out $f$ and $c$ (in a Bayesian sense) is equivalent to adjusting $f$ and $c$ at each step

Table S1: Number of independent simulations $N_{\text{runs}}$, force constant $k_c$ and simulation time per replica $T_{\text{sim}}$ (after removing 8 ns for equlibration) for a runs with 1, 2, 4 10 and 20 replicas

| **# of replicas** | $k_c$ | $N_{\text{runs}}$ | $T_{\text{sim}}$ |
|---|---|---|---|
| **1** | 12 | 20 | 450 |
| **2** | 5 | 8 | 460 |
| **4** | 3 | 4 | 290 |
| **10** | 1 and 0.5 | 6 | 340 |
| **20** | 0.5 | 4 | 190 |

to the value that leads to the smallest biasing energy. Without adjusting the constant $c$, we did not achieve good agreement between calculated and experimental curve at wider angles, possibly owning to a small buffer subtraction uncertainty. Critically, the adjusted value of $f$ and $c$ values were nearly identical among all SAXS-driven MD simulations, suggesting that $f$ and $c$ were not overfitted. After the SAXS-driven simulations had finished, we computed the SAXS curve from the entire refined ensemble. To compare this ensemble-averaged calculated curve with the experimental curve, a constant set of $f$ and $c$ was applied throughout this study ($f = 7.62433 \times 10^7$, $c = -22883e^2$), motivated from the fact that the adjusted $f$ and $c$ were highly similar in all SAXS-driven simulations.

**Calculations of semi-axes and shape averages.** In our previous work, semi-axes of the hydrophobic core ($a$, $b$, $c$) were calculated from the density profiles of the hydrophobic core.[5] This procedure requires averaging over a few nanoseconds for obtaining reasonable estimates of $a$, $b$, $c$. However, for obtaining distributions along $a$, $b$, $c$ as derived here, instantaneous values for $a$, $b$, $c$ are required. Hence, we estimated the instantaneous $a$, $b$, $c$ from the instantaneous moments of inertia (MOI) of the micelle, while assuming an ellipsoidal shape. We found that $a$, $b$, $c$ calculated from the MOI are systematically larger by $\sim 0.2$ nm as compared to the values calculated from the density profiles reported previously.[5]

The MOI of a tri-axial ellipsoid of mass $m$ and semi-axes $a$, $b$ and $c$ are given as:

$$\begin{aligned} I_a &= \frac{m}{5}(b^2 + c^2) \\ I_b &= \frac{m}{5}(a^2 + c^2) \\ I_c &= \frac{m}{5}(a^2 + b^2) \end{aligned} \tag{S6}$$

From the MOI, semi-axes of the tri-axial ellipsoid can be calculated as:

$$\begin{aligned} a &= \left(\frac{5}{2m}(I_b + I_c - I_a)\right)^{1/2} \\ b &= \left(\frac{5}{2m}(I_a + I_c - I_b)\right)^{1/2} \\ c &= \left(\frac{5}{2m}(I_a + I_b - I_c)\right)^{1/2} \end{aligned} \tag{S7}$$

To compute semi-axes from MD simulations, we first computed the three MOI of the micelle core every 10 ps using the GROMACS tool `gmx principal`. Assuming that the density of the core can be approximated by the ellipsoid of uniform density, the semi-axes were computed using Eqs. S7, and the distributions were computed from the refined ensemble.

**Modeling shape fluctuations.** For modeling shape fluctuations, samples of $a$, $b$, $c$, were drawn from the respective distributions. Here, we drew the samples from the one-dimensional distributions of $a$, $b$, $c$, thereby neglecting correlations between the semi-axes. To ensure that all semi-axes samples model a micelle with a constant volume, we normalized each drawn set $(a, b, c)$ via

$$\begin{aligned} \tilde{a}_i &= a_i \left(\frac{\langle V^{\text{core}}\rangle}{V_i^{\text{core}}}\right)^{1/3} \\ \tilde{b}_i &= b_i \left(\frac{\langle V^{\text{core}}\rangle}{V_i^{\text{core}}}\right)^{1/3} \\ \tilde{c}_i &= c_i \left(\frac{\langle V^{\text{core}}\rangle}{V_i^{\text{core}}}\right)^{1/3} \end{aligned} \tag{S8}$$

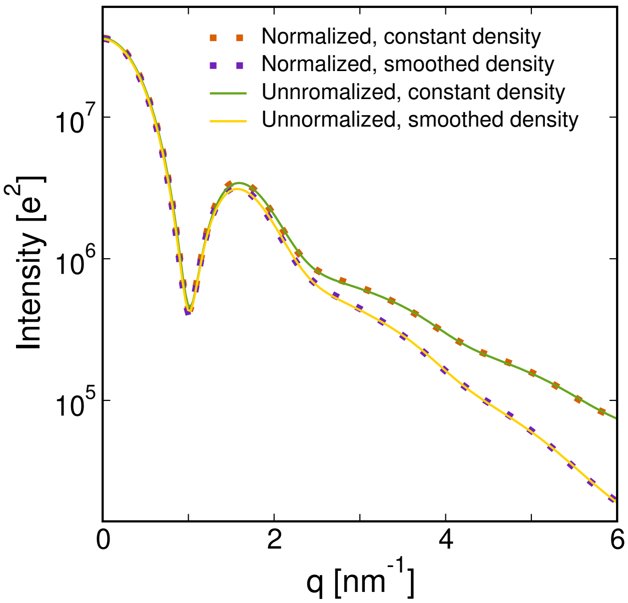


Figure S1: SAXS curves computed from the two-component tri-axial model. The density profiles were taken as piecewise constant (green line) or smoothed with a Gaussian filter (yellow curve). Samples of normalized and not normalised sets of semi-axes are taken from a 100 ns 4-replica simulation.

where $V_i^{\text{core}} = 4\pi abc/3$ is the volume of the ellipsoid with semi-axes $a, b, c$. The mean volume was taken as $\langle V^{\text{core}} \rangle = 4\pi \langle a \rangle \langle b \rangle \langle c \rangle /3$, where $\langle \cdot \rangle$ denotes the average over the respective refined ensemble. $\langle V^{\text{core}} \rangle$ differed only marginally from $4\pi \langle abc \rangle /3$. Subsequently, the SAXS curve was computed using the analytic model of the tri-axial ellipsoid using the normalized set of semi-axes $(\tilde{a}, \tilde{b}, \tilde{c})$, and the SAXS curves were averaged. Notably, the correction of the semi-axes via Eqs. S8 had only a marginal effect on the SAXS curves (Fig. S1). Errors were computed as 1SEM using only independent multi-replica simulations as independent data points, providing a conservative error estimate.

### Notes on force field imperfections

During all ten-replica simulations, the micelle in one of the replicas adopted an unexpected horseshoe shape, leading to slightly larger value for the large semi-axes $a$ semi-axis (Fig. S2). Consistent with this observation, the micelle in two of 20 replicas adopted a horseshoe shape during 20-replica simulations. With fewer replicas, no such horseshoe shapes were observed. Although we can not strictly exclude that such shapes occasionally exist under experimental conditions, we speculated that either (i) unknown systematic errors in the data, (ii) overly restrained ensembles, or (iii) force field imperfections may be responsible for such occasional shapes.

To shed more light on these observation, additional test simulations were carried out. First, motivated by the fact that the experimental curve exhibits some uncertainty around the pronounced minimum ($q \approx 1\,\mathrm{nm}^{-1}$), we performed two addition sets of refinement simulations using target SAXS curves, whose error was increased by factors of 3 or 5 in the $q$-region between $0.85\,\mathrm{nm}^{-1}$ and $1.25\,\mathrm{nm}^{-1}$, leading to strongly reduced weights in this $q$-region. However, also in these additional simulations, the horseshoe shapes were reproduced in one out of ten replicas, suggesting that the relatively high uncertainty at the $q \approx 1\,\mathrm{nm}^{-1}$ region does not cause the horseshoe shapes. Second, we performed series of test simulations with weaker coupling to the experimental curve, by reducing the force constant ($k_c$). Only with very low $k_c = 0.1$, the horseshoe shape vanished, but now the agreement to the experimental data was significantly reduced.

Taken together, it seems unlikely that systematic experimental errors induced the horseshoe shapes, or that the simulations were overly restrained to the data. Instead, we hypothesise that, with increasing number of replicas, and hence an increasing number of degrees of the freedom, ensemble refinement becomes more sensitive to force-field imperfections. Hence, ensemble refinement, as conducted here, is also a starting point for future developments of soft matter force fields.

To exclude that the occasional horseshoe shape influences the conclusions of this manuscript, we report results from four-replica simulation, in which no horseshoe shapes were observed, along with results from ten-replica simulations. The distributions of $a$, $b$, $c$ were similar in four- and ten-replica simulations (except for contributions from the horseshoe-shaped micelles, Fig. S2), and the mean values of $a$, $b$, $c$ are nearly identical. This suggests that the key conclusions were not affected by the force field imperfections.

#### Note on solvent simulations

All SAXS-driven simulations were conducted in 150 mM NaCl solution, while the SAXS experiment was performed in pure water solvent. To exclude that the details of the solvent influence the conclusions of this study, we repeated the key simulations with the DDM micelle in pure-water solvent. In addition, to exclude that the solvent density affects the calculations with the two-component model, we repeated the fits with the two-component model assuming a solvent density of $334.7\,\mathrm{e\,nm^{-3}}$ to match the electron density of water at 15°C.[1] The fitted parameters, as shown in Table S2, are nearly identical to the results found with the 150 mM NaCl solution (Table 1).

Table S2: Average semi-axes calculated from multi-replica SAXS-driven MD simulations (top rows) and from fitting two-component ellipsoid models (bottom rows), here assuming pure-water solvent. Errors of $a$, $b$, $c$ were computed using block averaging with 4 ns blocks or estimated as 1 SEM of averages between independent runs, and errors were typically smaller than 0.03 nm.

| **All-atom MD # of replicas** | **a [nm]** | **b [nm]** | **c [nm]** | **t [nm]** |
|---|---|---|---|---|
| 1 | 3.30 | 2.30 | 1.66 | |
| 4 | 3.19 | 2.38 | 1.68 | |
| 8 | 3.19 | 2.43 | 1.66 | |
| 24 | 3.19 | 2.43 | 1.69 | |
| **Two-component model** | | | | |
| prolate (unphysical) | 3.40 | 1.99 | 1.99 | 0.55 |
| oblate | 2.85 | 2.85 | 1.60 | 0.53 |
| tri-axial | 3.22 | 2.48 | 1.66 | 0.53 |

#### Supporting figures

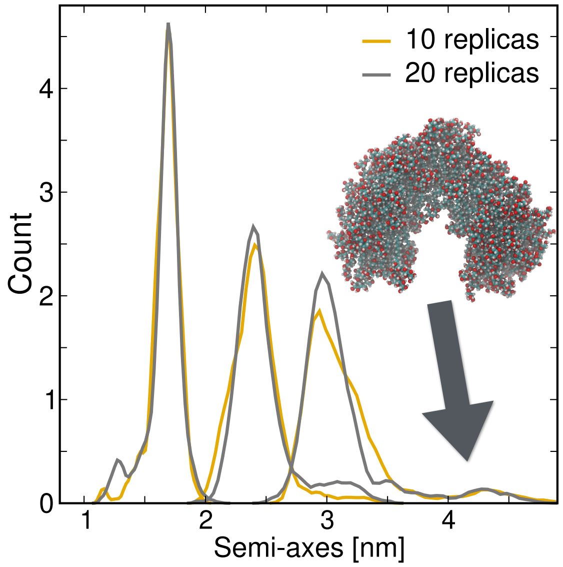


Figure S2: Distributions of semi-axes in simulations with 10 or 20 replicas, see legend. During these simulations, the micelle adopted in one of ten replicas a horseshoe shape. See SI Text for further discussion. Further, the agreement among the 10- and 20-replica simulations suggests that 10 replica are sufficient to achieve a minimally biased ensemble.

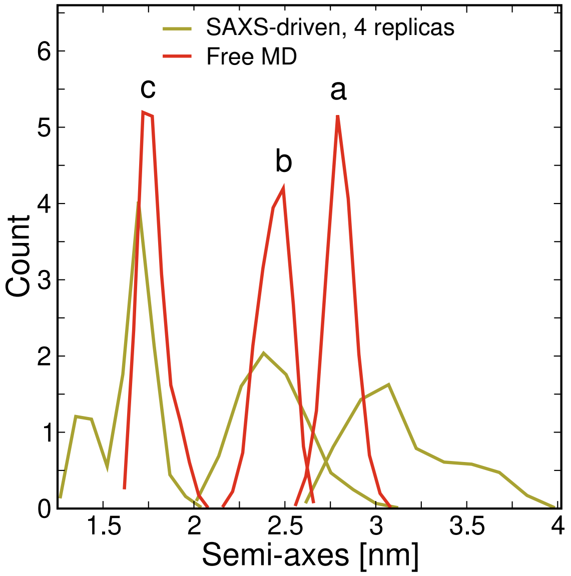


Figure S3: Distributions of semi-axes, calculated from a four-replica SAXS driven simulation (yellow) or free a MD simulation (red). Evidently, the three axes are slightly more similar in free as compared to SAXs-driven simulations, corresponding to a slightly too spherical shape.[5]

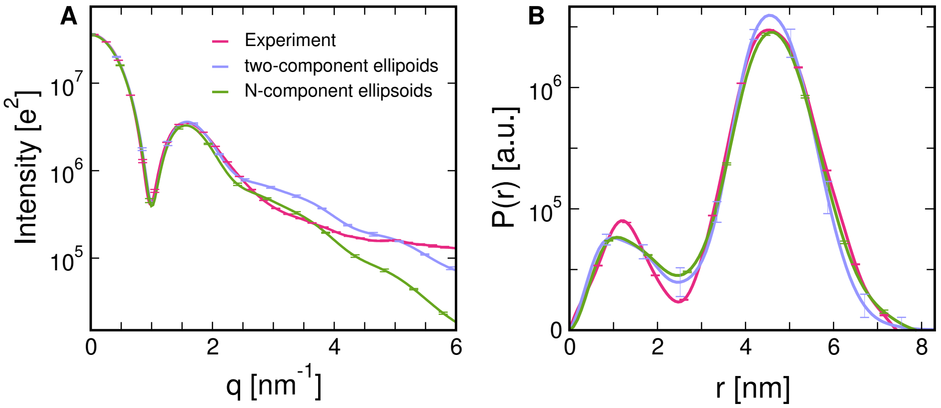


Figure S4: (A) SAXS curve computed as an average of sets of two-component tri-axial ellipsoids (blue), or of $N$-component tri-axial ellipsoids with smoothed electron densities around the headgroup region (green). Both models exhibit poor agreement with experimental data (red) at wider angles, suggesting that modeling of atomic details is mandatory at wide angles. Here, the sets of semi-axes were taken from 10-replica MD simulations. Curves calculated with semi-axes from four-replica simulations are nearly identical and shown in Fig. 3.

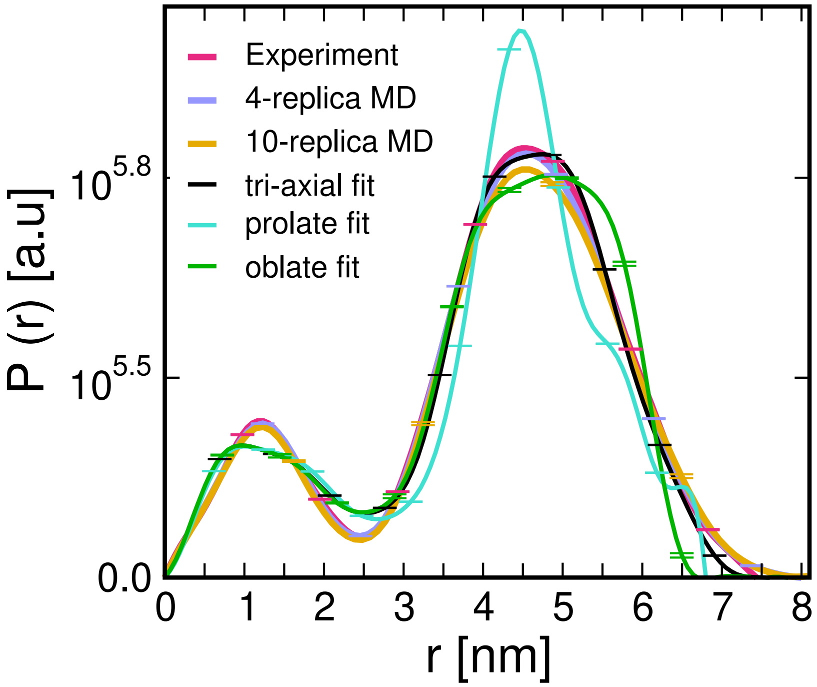


Figure S5: Pair-distance distribution functions $P(r)$ with the representative errors, obtained from the experimental SAXS curve, from MD-derived SAXS curves, or from curves of fitted oblate, prolate and tri-axial models (see legend for color code). The $P(r)$ curves were computed using GNOM[2] with default settings. The parameter for the maximum diameter ($D_{\max}$) of the particle was taken from the MD frames or from the fitted structural models. $P(r)$ from experiment and from four-replica MD agree favorably. The small deviations between experiment and 10-replica MD at $r \sim 4.5$ nm is a consequence of the horseshoe-shaped micelle in one of ten replicas, see SI Text for discussion and Fig. S2. $P(r)$ from the fitted tri-axial ellipsoid give a reasonable agreement to experiment, while $P(r)$ from prolate/oblate fits reveals major discrepancies to the experiment.

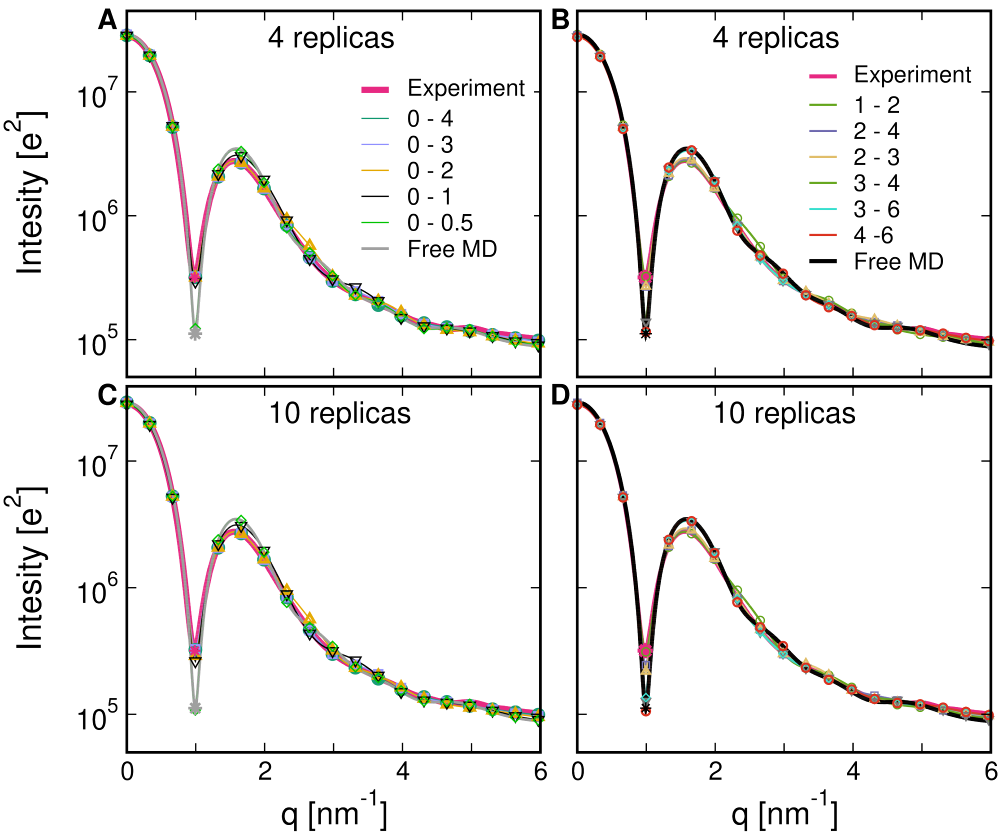


Figure S6: SAXS curves computed from four-replica SAXS-driven simulations (A and B) and 10-replica SAXS-driven simulations (C and D), using only $q$-intervals of the experimental curve as a target. The applied $q$-ranges are indicated in the legends. Representative symbols with the same colors codes are shown to guide the eye.
